# Microvascular immunity is organ-specific and concealed in peripheral blood

**DOI:** 10.1101/2024.04.02.584356

**Authors:** Rebecca Rixen, Paula Schütz, Carolin Walter, Birte Hüchtmann, Veerle Van Marck, Barbara Heitplatz, Julian Varghese, Hermann Pavenstädt, Konrad Buscher

## Abstract

Blood tests are a common method for diagnosing and monitoring various health conditions. Nevertheless, the extent to which phlebotomy can offer insights into immune and organ dysfunction remains uncertain. Here, we conducted a comprehensive analysis of blood-borne leukocytes in the microvasculature of different mouse organs and compared it to peripheral blood and parenchymal samples. We observed that microvascular immune cells outnumber tissue-resident counterparts in the kidney, liver and lung. Classical monocytes and lymphocytes are diminished while nonclassical and SSC-high monocytes are enriched compared to blood. Utilizing single-cell sequencing, we identified specific cell populations up to 100-fold expanded in the kidney vasculature including macrophages, plasmacytoid dendritic cells, B cells, and innate lymphoid cells type 2. Microvascular enrichment could trigger a local phenotype switch as shown in glomerulus-restricted B cells. Peritonitis and acute kidney injury (AKI) elicited a multifaceted and systemic response of microvascular leukocytes. It involved remote organ effects, such as a 16-fold increase of leukocytes in the splenic circulation or 64-fold increase of SSC-high monocytes in the liver circulation that was not detectable in the peripheral blood or the tissue. Following full recovery from AKI, persistent and complex changes were observed predominantly in the renal vasculature, while most leukocytes in the peripheral blood had already returned to baseline levels. Collectively, our findings suggest a paradigm of organ- and disease-specific microvascular immunity that largely eludes conventional blood and tissue analysis.

## Introduction

A comprehensive understanding of an organism’s immune state holds profound clinical relevance ^1–3^. Venous blood samples are routinely employed to explore cell, protein and molecule concentrations. However, immunity is significantly shaped in lymphoid organs and local tissue. From an immunological perspective, the blood circulation is mostly perceived as a conduit, facilitating the transit of immune cells, mediators and metabolites to virtually every corner of the body. Immune surveillance and the swift mobilization of effector cells heavily rely on the seamless connectivity between the local and systemic environments. Therefore, the extent to which blood biomarkers can capture the complexities of immune dynamics in health and disease remains an open question ^4,5^.

Organs harbor resident leukocytes that play a pivotal role in immunity. They encompass a spectrum of cells critical in innate and adaptive immune responses, including mononuclear phagocytes (such as monocytes, macrophages, and dendritic cells (DC)), natural killer (NK) cells, lymphocytes, and innate lymphoid cells (ILCs) ^6^. While some of these cell types remain sessile within the tissue, others engage in continuous recirculation through the lymphatic and circulatory systems ^6^. The microcirculation acts as an interface between the tissues and the bloodstream, facilitating the extravasation of blood-borne cells into the parenchyma during homeostasis and inflammation. Beyond their transit function, endothelial cells also enable intravascular immune surveillance ^7^. Some immune cells have been identified as preferentially residing within the vasculature. Non-classical monocytes and CD8 T lymphocytes, for instance, actively “patrol” the endothelium to detect cues of injury ^8^. In atherosclerosis, these monocytes accumulate even in high-shear arteries to safeguard the endothelial barrier ^9^. Kupffer cells, located intravascularly within the liver sinusoids, screen for blood-borne antigens and bacteria. In the lung, intravascular macrophages, marginating B cells, and iNKT cells have been documented ^10–12^.

Together, the microcirculation represents a topological niche of high significance for our understanding of immunity. Nevertheless, it frequently eludes direct scrutiny, and microvascular immune cells at the blood-tissue interface have been rarely studied. Here, we implemented a protocol to label microvascular leukocytes in different mouse organs and comprehensively analyzed their numbers, phenotypes and inflammatory changes in direct comparison to matched peripheral blood samples.

## Methods

### Antibodies and reagents

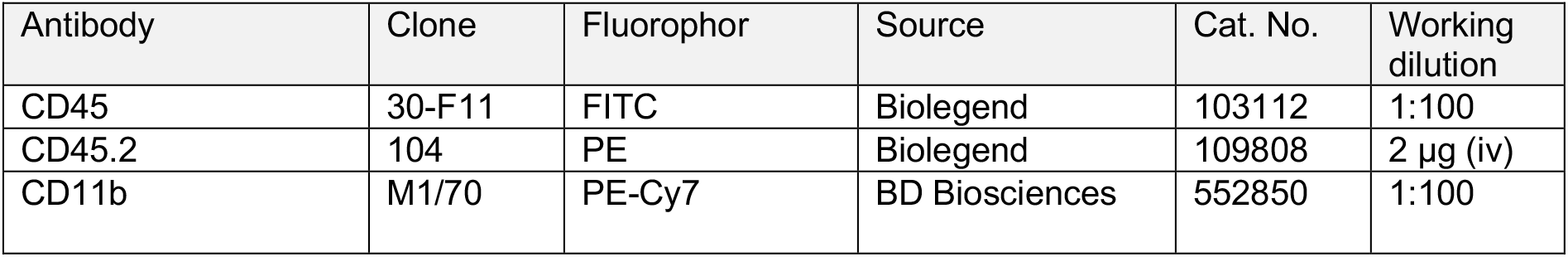
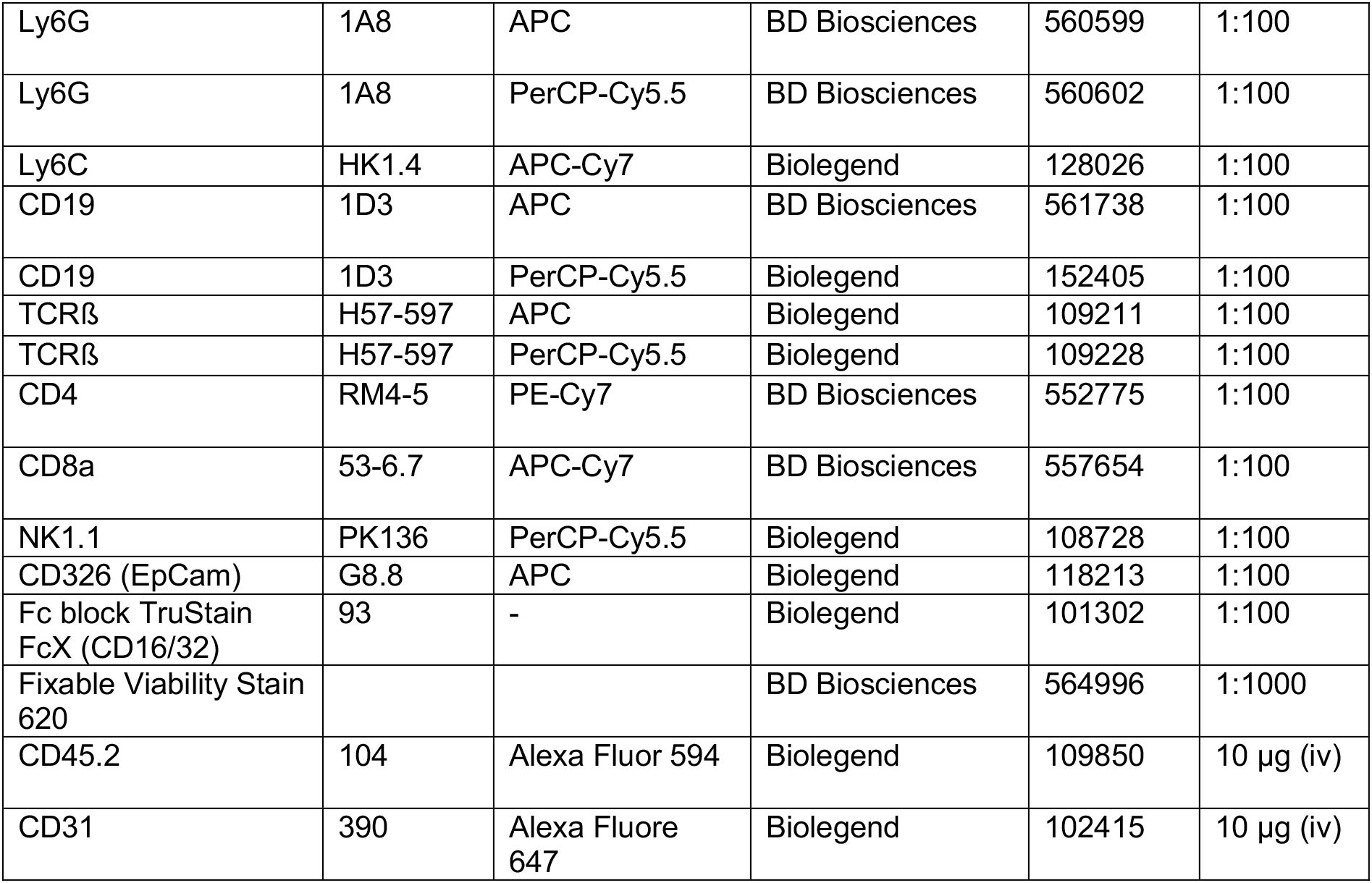

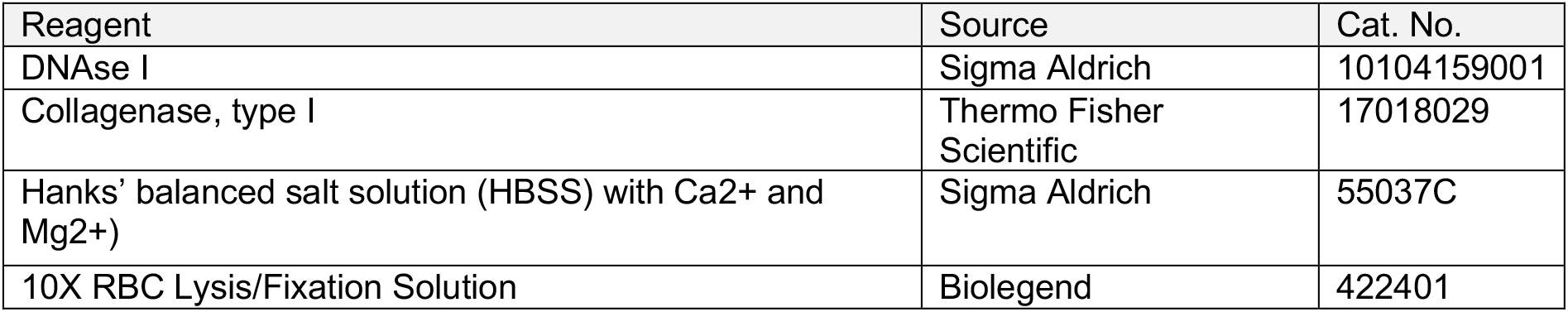

| Reagent | Source | Cat. No. |
| --- | --- | --- |
| DNAse I | Sigma Aldrich | 10104159001 |
| Collagenase, type I | Thermo Fisher Scientific | 17018029 |
| Hanks' balanced salt solution (HBSS) with Ca<sup>2+</sup> and Mg<sup>2+</sup>) | Sigma Aldrich | 55037C |
| 10X RBC Lysis/Fixation Solution | Biolegend | 422401 |

### Mice and disease models

All animal experiments were approved by the authorities (Landesamt für Natur, Umwelt und Verbraucherschutz, approval number: 81-02.04.2019.A253) and performed in accordance with the animal protection guidelines of Germany. C57BL/6 mice were purchased by Charles River and bred in house. Mouse models of peritonitis and bilateral renal ischemia/reperfusion injury (IRI) were studied at day 1 (INF/AKI) and day 12 (INF-reg/AKI-reg) post injury. The surgical procedure was performed as follows: Buprenorphine (0.1 mg/kg) was administered subcutaneously to 8-20 week-old male mice 20 min prior to surgery. Mice were anesthetized with a 1,5-2% isoflurane/oxygen mixture, placed on a thermoregulated pad to maintain body temperature, and a midline abdominal incision was made. For AKI-IRI induction, both renal pedicles were carefully dissected and clamped with an atraumatic vascular clamp for 35 min.

For inducing semi-sterile peritonitis (INF, INF-reg), the cavity was left open for 35 min and the gut was mobilized similar to AKI-IRI. Animals received buprenorphine via the drinking water (0.009 mg/ml) up to 3 days post-surgery.

### Intravascular stainings

Due to reported detrimental effects on the tissue integrity, we did not flush the vasculature prior to organ collection ^13^. Hence, blood-borne non-interacting leukocytes are present in the explanted organ. To account for this effect, we used peripheral blood as a reference. We opted for right ventricular blood as it closely resembles the composition of blood in larger peripheral veins typically accessed for venipuncture in human patients. For intravascular leukocyte stainings, a Phycoerythrin (PE)-labeled rat anti-mouse CD45.2 antibody (2 µg) was injected intravenously. After 5 minutes of incubation, blood was collected by right cardiac puncture and organs were harvested without vascular perfusion. Isotype controls are shown in supplemental figure 1. To exclude the staining of the extravascular cells using this protocol, the epithelial cell antibody anti-EpCam (CD326, APC-labeled) was injected (supplemental figure 1). For imaging experiments, 10 µg of fluorescently-labeled antibodies (CD45.2-AF594, CD31-AF647) were injected intravenously. Validation experiments were performed for untreated mice, AKI and AKI-reg (supplemental figure 1).

### Flow cytometry and cell sorting

Heparinized blood was incubated with Fc Block (Biolegend) for 10 min on ice followed by antibody staining for 30 min. Erythrocytes (RBC) were lysed for 10 min at room temperature (RT) using 1x RBC Lysis/Fixation solution (Biolegend). Samples were washed with phosphate buffered saline (PBS) at 400x g for 5 min and cell pellets were resuspended in PBS with 1% fetal calf serum (FCS). Organs were minced into small pieces, and kidney, liver and lung were digested in 1 mL HBSS (with Ca/Mg) including DNAse I (10mg/mL) and collagenase I (100 U/µL) at 37°C and 350 rpm for 30 min (kidney) or 45 min (liver and lung). Addition of 2 ml PBS to the digestion buffer was followed by filtering suspensions through a 70 µm cell strainer. Samples were then centrifuged at 400g for 5 min and cell pellets were resuspended in PBS with 1% FCS. Cells were incubated with antibody mixtures (1:100) for 30 min on ice, followed by fixation with 4% PFA for 10 min. RBC lysis was also required for the spleen tissue. Cells were washed with 2 ml PBS (400g, 5 min) and resuspended in PBS with 1% FCS. All samples were measured using BD FACSCanto II (BD Biosciences). Data analysis was performed using FlowJo v10.9. software (BD Bioscience). Fluorescence-activated cell sorting was used to extract viable CD45+ leukocytes from the blood, kidney microcirculation (CD45+CD45.2^+^), and kidney tissue (CD45^+^CD45.2^-^) as described above using a BD FACS Aria II flow cytometer equipped with 11 detectors and FACSDiva software. Cells were sorted into tubes containing 200 µl PBS + 2% FCS on ice and washed with PBS once before further processing.

### Immunofluorescence microscopy

Freshly harvested tissue was cryopreserved in 10% sucrose solution overnight at 4°C, embedded in Tissue Tek O.C.T (Sakura Finetek), frozen in liquid nitrogen and then stored at -80°C. 7 µm sections were cut with a cryostat (Thermo Scientific, blade temperature: -25°C, specimen temperature: - 15°C) and mounted on superfrost microscope slides. Sections were fixed with 4% paraformaldehyde (PFA) for 10 min, washed three times 5 min with PBS. For ex vivo staining, sections were blocked with 10% goat serum/5% bovine serum albumin (BSA) in PBS for 2h at room temperature and then incubated with primary antibodies diluted in 5% goat serum/5% BSA in PBS overnight at 4°C in a humidity chamber. After three washing steps, secondary antibodies diluted in 5% goat serum/5% BSA in PBS were applied on the sections for 2h at room temperature. Cryosections were washed three times 5 min with PBS and were mounted with Fluoroshield with DAPI (Sigma Aldrich, F6057). Images were acquired using an Axio Imager M2 with Apotome 2.0 and 20x objective (Zeiss, Germany). For imaging of thick tissue sections, organs were fixed in 4% PFA overnight and were embedded using 3% low-melting agarose. 50-100 µm sections were cut with a vibratome (Leica, VT1000 S). For SDS delipidation, samples were incubated in a clearing solution (200 mM boric acid and 4% SDS, pH 8.5) at 70 °C at 500 rpm for 1 hour using a benchtop shaker. Free-floating sections were washed with PBS + 0,1 % Triton-X 100 (PBST) for 5 min at RT followed by incubation with primary conjugated antibodies in PBST overnight at 4°C. After a further washing step with PBST, sections were mounted with Fluoroshield with DAPI onto microscope slides for confocal imaging (Leica, SP8 confocal microscope, 40x water objective). ImageJ/Fiji (http://fiji.sc/) and Zen Software (Zeiss, Germany) were used to analyze images.

### Histology and scoring of kidney injury

Paraffin-embedded kidney sections were stained with hematoxylin and eosin (H&E) and periodic acid-Schiff (PAS) using standard protocols. Tubular injury based on casts, dilatation and damage was scored by two independent pathologists trained in nephropathology.

### Laboratory values

Heparinized cardiac blood was collected and centrifuged at 2000x g for 20 min. Plasma was transferred into cryotubes and stored at -20°C. Creatinine and blood urea nitrogen were measured by IDEXX Laboratories (Kornwestheim, Germany).

### Targeted RNA single cell library preparation and sequencing

For single-cell RNA sequencing, the BD Rhapsody Express System (BD Biosciences) was used according to the manufacturer’s protocols. Viable CD45+/CD45.2+ (microcirculation), CD45+/CD45.2- (tissue) and CD45.2+ peripheral blood leukocytes were FACS-sorted and labeled using the BD Rhapsody Single-Cell Multiplexing CTT Kit (BD Biosciences, 626545). On average, a total of 12.944 leukocytes were captured per mouse after cell sorting (5.868-21.577). Sample tags were incubated at room temperature (RT) for 20 minutes and washed three times. The pooled sample was resuspended in an ice-cold sample buffer (BD Biosciences, BD Rhapsody Cartridge Reagent Kit, 633731). Isolation of single cells was performed using the single cell capture and cDNA synthesis kit according to the manufacturer’s protocol. Briefly, the samples were loaded onto the primed nanowell cartridge (BD Biosciences, 633733), incubated at room temperature for 15 min followed by cell capture beads and further incubation of 3 min at room temperature. Cells were lysed followed by bead retrieval and washing. Reverse transcription was performed for cDNA synthesis following Exonuclease I treatment (BD Biosciences, 633773).

Libraries were prepared using the BD Rhapsody targeted mRNA and Abseq amplification kit (BD Biosciences, 633774). Targeted amplification of cDNA was performed using a mouse immune response panel (397 genes, BD Biosciences, 633753) and a custom additional panel of 99 genes (covering leukocyte lineage- and kidney cell-specific genes, supplemental table 1) by PCR (11-15 cycles). For double-sided DNA size selection, Agencourt AMPure XP magnetic beads (Beckman Coulter, A63880) were used to separate sample tag PCR products from mRNA target PCR products. Further amplification of the sample tag and the mRNA-targeted PCR products was performed by PCR (10 cycles). The PCR products were then purified using Agencourt AMPure XP magnetic beads. The concentration of each sample was determined using a Qubit fluorometer and the Qubit dsDNA HS Assay Kit (Thermo Fisher Scientific, Q32851). To prepare the final libraries, the purified PCR products were indexed by PCR (6-8 cycles). The index PCR products were purified using Agencourt AMPure XP magnetic beads. Quality control was performed by estimating the concentration and measuring the average fragment size of the mRNA target library and sample tag library using the Agilent Tape Station with the High Sensitivity D1000 ScreenTape (Agilent, 5067-5584). The final libraries were diluted to a concentration of 4 nM and multiplexed for Novaseq paired-end sequencing (150 bp) including 20% PhiX spike-in. The sequencing depth was calculated with 600 and 4000 reads/cell for the sample tag and the mRNA library, respectively. Sequencing was performed using the Illumina NovaSeq 6000 sequencer.

### Sequencing data analysis

Demultiplexing and preprocessing of the sequencing data was conducted with the BD Rhapsody sequence analysis pipeline v.1.11 from SevenBridges according to the manufacturer’s protocol. Sequences were aligned against a targeted genome panel based on the murine reference mm10. All error-corrected scRNA count matrices were imported into R v4.0.5 and Seurat v4.0.5 ^14^. For each sample, only cells with at least five features were considered, and rare features that were present in less than 10 cells per sample were discarded. To remove potential doublets or multiplets from the analysis, cells with a very high RNA count value were filtered, with thresholds ranging from 2.000 to 10.000 depending on the sample’s nCount_RNA value distribution. Subsequently, Seurat’s SCTransform routine was used to normalize and integrate the scRNA samples for each data set, while PCAs and UMAPs were employed for dimension reduction. Finally, clusterings for each data set were created with Seurat’s FindClusters function, using a resolution of 0.5 and default values otherwise. Additionally, all identified B cells were extracted and reclustered with a resolution parameter of 0.2 to allow for a more detailed view into this cell type of interest. For all data sets and subsets, clusters were annotated using a combined approach of marker gene visualization and differential gene expression. Target genes were selected and expression patterns were visualized with Seurat’s FeaturePlot functionality. The package’s FindMarkers routine was chosen to identify differentially expressed genes per cluster using the MAST algorithm ^15^. Pseudotime and trajectory analysis was conducted with the R/Bioconductor package Slingshot v1.8 ^16^. Briefly, Seurat cell embeddings were imported and lineages were calculated based on the chosen start cluster of interest, the main B cell cluster of the peripheral blood. Pseudotime and tree visualizations were then realized with Slingshot’s plot functions. For receptor ligand analysis, we merged our sequencing data set (as shown in figure 5) with the mouse endothelial data E-MTAB-8145 by filtering for common genes and applying the integration algorithm Harmony ^17^. E-MTAB-8145 includes untreated murine renal endothelial cells from glomerular, cortical and medullary kidney compartments. CellPhoneDB’s statistical-analysis method (v2.1.5) was applied to identify potential receptor-ligand interactions between leukocyte and endothelial clusters ^18^. The minimum interaction score was set to 0.037 and the maximum interaction p value to 0.05.

### Data availability

The sequencing data was uploaded in the repository GSE252496.

### Statistics

Statistical tests are indicated in the figure legend. *** p < 0.001, ** p < 0.01, * p < 0.05

## Results

We adopted an intravascular staining protocol with intravenous injection of an anti-CD45.2 leukocyte antibody prior to euthanasia ^13^. A circulation time of 5 minutes did not lead to extravascular staining as confirmed by using the epithelial (= extravascular) marker EpCAM (supplemental figure 1). Ex vivo staining of CD45 allowed us to separate microvascular (CD45+ CD45.2+) from extravascular (CD45+ CD45.2-) leukocytes (figure 1a, supplemental figure 1). We did not perfuse the organs in situ to avoid losing marginating (temporarily adhered) cells in the microcirculation and to prevent possible disruption of the organ architecture^13^.

**Figure 1:**
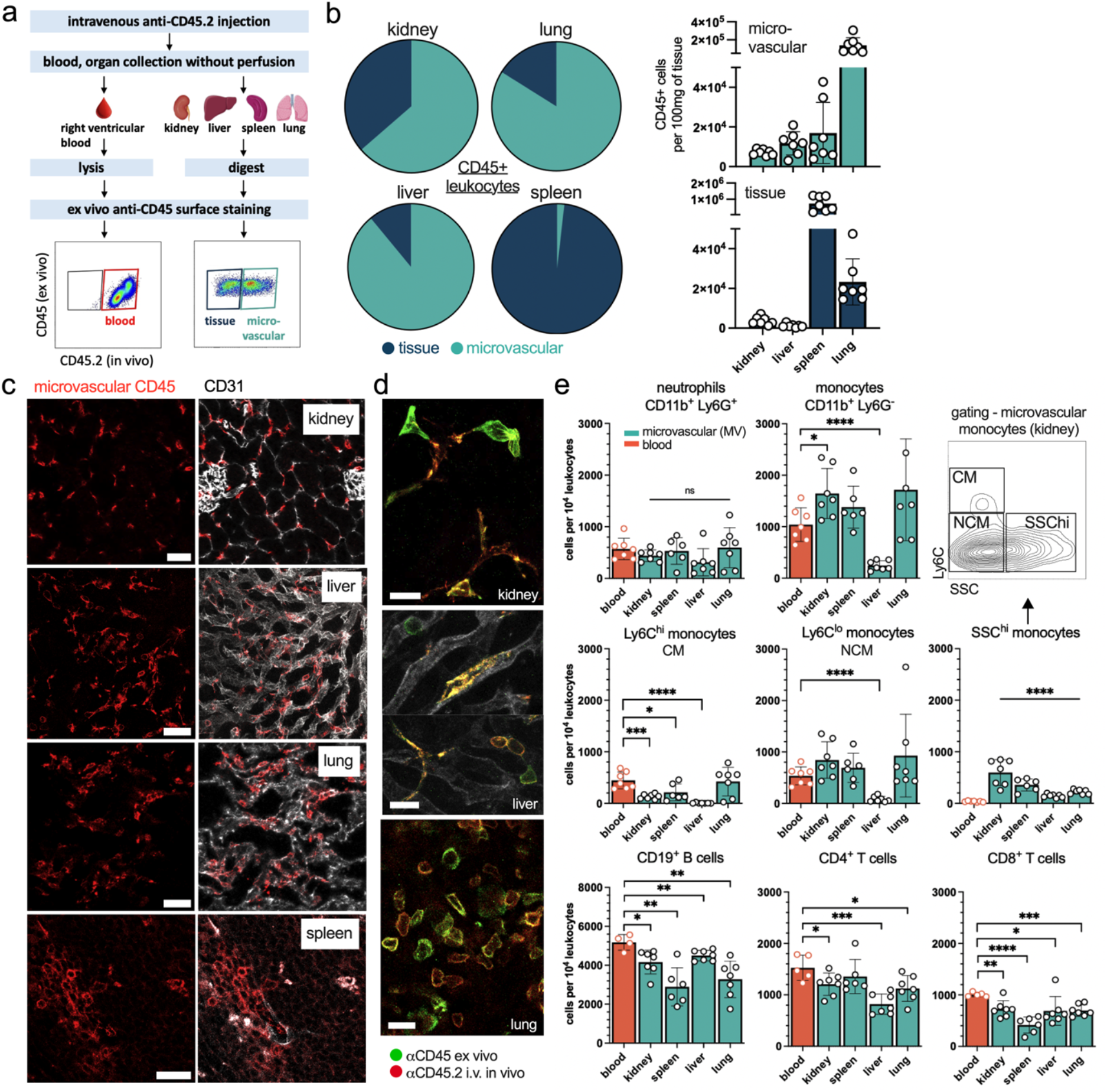
Microvascular leukocytes are abundant, organ-specific and differ from peripheral blood. a) Experimental setup to detect microvascular and tissue-resident leukocytes in mice. A matched sample of right ventricular blood allows for direct comparison. b) The total numbers of leukocytes (right panels) and the relation of microvascular and tissue-resident leukocytes (pie charts) in different healthy mouse organs. c) The injection of an anti-CD45 antibody labels intravascular leukocytes in healthy organs. Scale bar 100 μm. d) The combination of in vivo and ex vivo anti-CD45 staining highlights vascular and tissue-resident leukocytes. Intraluminal cell extensions (double positive) of perivascular cells (green) are predominantly found in the kidney. Scale bar 20 μm. e) Flow cytometry analysis of different intravascular leukocyte subsets in comparison to peripheral blood (red bar). Unpaired t-test of blood versus each organ.

In the kidney, lung and liver, vascular leukocytes largely outnumbered parenchymal leukocytes to varying degrees (figure 1b). The high abundance of microvascular leukocytes was also evident in immunofluorescence stainings (figure 1c, supplemental figure 2). The combination of in vivo and ex vivo CD45 staining highlighted that some leukocytes are located extravascularly with intraluminal extensions, predominantly in the kidney (figure 1d). We assessed different leukocyte lineages in these organs by flow cytometry (gating strategy in supplemental figure 1). Microvascular cell populations were compared to a blood sample that was obtained at the time of organ harvest by right-ventricular puncture. There was no difference in neutrophil numbers (Lin^-^ CD11b^+^ Ly6G^+^) between the blood and any organ vasculature (figure 1e). In contrast, monocyte numbers (Lin^-^ CD11b^+^ Ly6G^-^ side scatter (SSC)^lo^) significantly differed. Kidney, spleen and lung showed increased microvascular monocytes, which was mainly caused by the nonclassical (Ly6C^lo^ SSC^lo^) subset. In contrast, classical monocytes (Ly6C^hi^ SSC^lo^) are mostly reduced in the microcirculation. A noticeable exception was the liver vasculature where SSC^lo^ monocytes are largely depleted. SSC high monocytes are more granular and represent a macrophage-like phenotype (figure 1e). They were detectable in all organs, predominantly in the kidney, but not in blood. B cells (Lin^-^ CD19^+^) and T cells (Lin^-^ CD3^+^ CD4^+^ or CD8^+^) were reduced in the microcirculation of most organs compared to blood (figure 1e).

To detect unsupervised phenotypes and transcriptional states of microvascular leukocytes in comparison to the peripheral blood we performed single cell RNA sequencing. Leukocytes in the renal tissue, the renal microvasculature (MV) and in the blood of one mouse were sorted, multiplexed and processed together on the same microwell device for library preparation. Using 4 healthy mice, 33.474 leukocytes were sequenced (13.828, 13.408 and 6.238 in blood, microcirculation and tissue, respectively), resulting in 27 CD45^+^ leukocyte clusters (figure 2a, supplemental figure 3, supplemental table 1).

**Figure 2:**
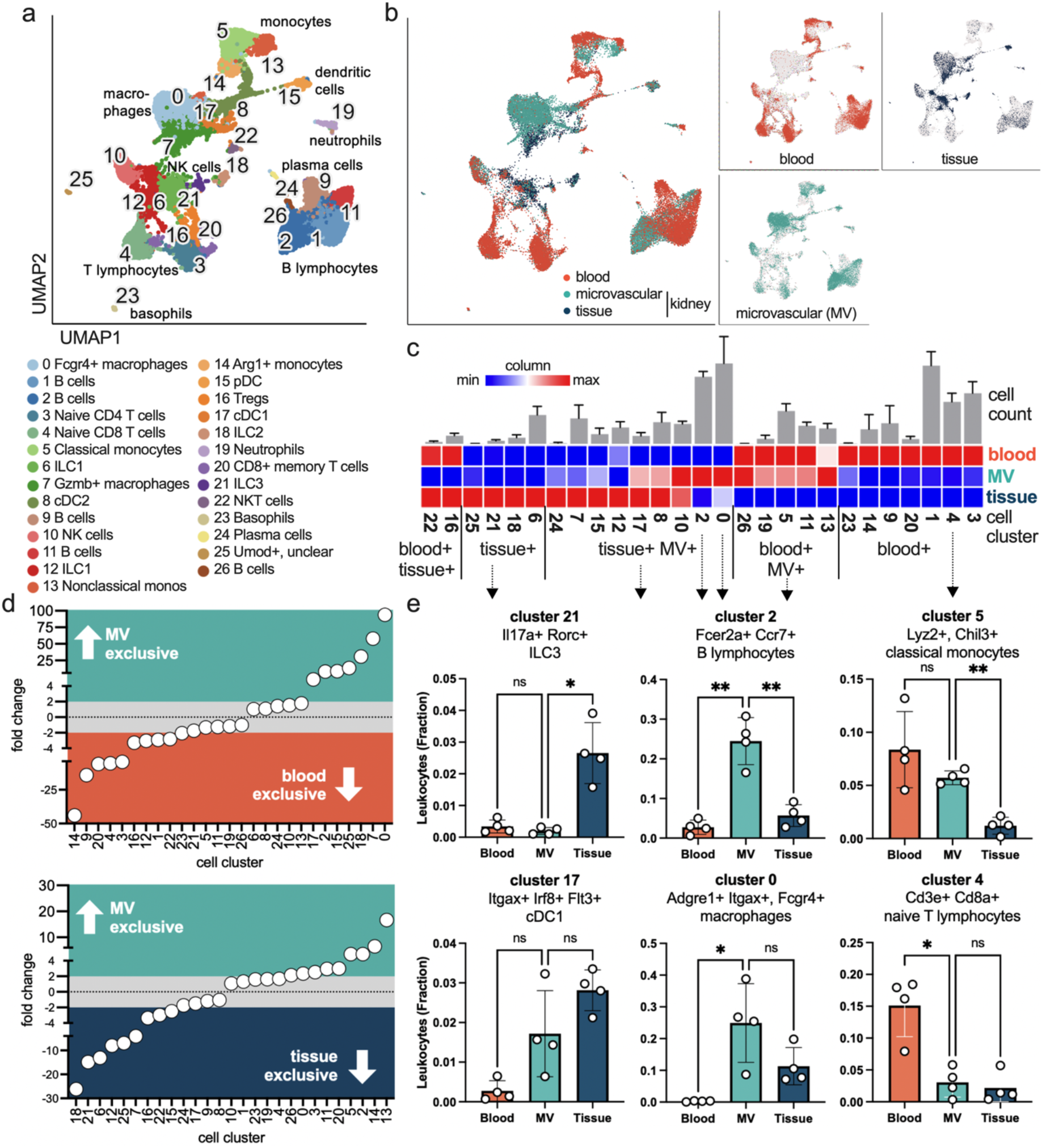
Unsupervised phenotyping of microvascular immune populations in the healthy mouse kidney. a) Single cell sequencing of leukocytes sorted from blood, the renal microcirculation and the renal tissue. UMAP visualization and annotation of 26 leukocyte subtypes. n=4 independent experiments. b) Demultiplexing of all leukocytes. c) Abundance of cell cluster specific leukocytes in the three compartments. Blue is the lowest number and red the highest number for each column. The top bar graphs represent the total cell count of each cluster as the mean +/-S.D. of n=4 experiments. d) Ranked fold change of microvascular versus peripheral blood (top), and microvascular versus tissue-resident (bottom) leukocytes for each cell cluster. e) Individual cell cluster depicted with the total number of leukocytes in the three compartments as fraction of all detected leukocytes. One-way Anova with Dunnett’s multiple comparison. MV = microvascular.

Demultiplexing assigned each cell to its origin (tissue vs. renal microvasculature vs. blood), demonstrating a cell type and compartment specific distribution (figure 2b-e, supplemental figure 3, supplemental table 1). As expected, the tissue-resident populations of innate lymphoid cells (ILC) types 1-3 (clusters 6, 12, 18, 21) had a strong tissue signature (figure 2d,e; supplemental figure 3). Interestingly, a number of leukocytes were preferentially found in peripheral blood mostly sparing the renal circulation (cluster 4, figure 2c-e). This included lymphocytes, confirming our flow cytometry data. Vice versa, seven cell types preferentially populated the renal circulation and are low or absent in the blood (figure 2c,d). This included cluster 0 as highly abundant Adgre1^+^ Itgax^+^ Fcgr4^+^ macrophages. They were almost 100-fold enriched in the renal vasculature compared to blood, and low in the tissue (figure 2c,d,e). A similar pattern (5-to 50-fold enriched) was detected in six other leukocyte types (cluster 2,7,15,17,18,25) including Gzmb^+^ macrophages, plasmacytoid dendritic cells (DC), conventional DCs type 1, ILC type 2, and B cells (figure 2c,d). Cluster 18 as ILC type 2 (Gata3^+^ IL13^+^) was assigned predominantly to the tissue but was also detectable in the microvasculature (20-fold lower but still 20-fold higher compared to blood), suggesting to a unique topology at the kidney-blood interface.

Cluster 2 was exclusively present in the renal vasculature, and absent in both blood and kidney tissue (figure 2c,d,e), pointing to a specialized endovascular B cell subset. Reclustering of all B cells showed a gradient between the blood and microcirculation (figure 3a,b). Of all renal B cells, 86% were microvascular and 14% were tissue-resident. Subclusters 6, 4 and 7 were 5-, 12- and 15-fold increased in the renal vasculature compared to blood, respectively (figure 3c). Slingshot, a lineage inference tool, identified a developmental trajectory from the main blood-borne B cell cluster 0 to the microvascular clusters (figure 3d). We found the transcription factor Early Growth Response-1 (Egr1) and Interferon Regulatory Factor 4 (Irf4) as main driver genes underlying this phenotype switch. The transcriptional shift included differentially regulated gene pathways in microvascular B cells including MAPK kinase cascade and interleukin 1 production (figure 3e). Using immunofluorescence imaging with intravascular staining of CD19, renal B cells were found almost exclusively in glomerular capillaries whereas tubular and medullary regions were largely depleted (figure 3f). These data suggest an endovascular B cell phenotype switch elicited by glomerular capillaries.

**Figure 3:**
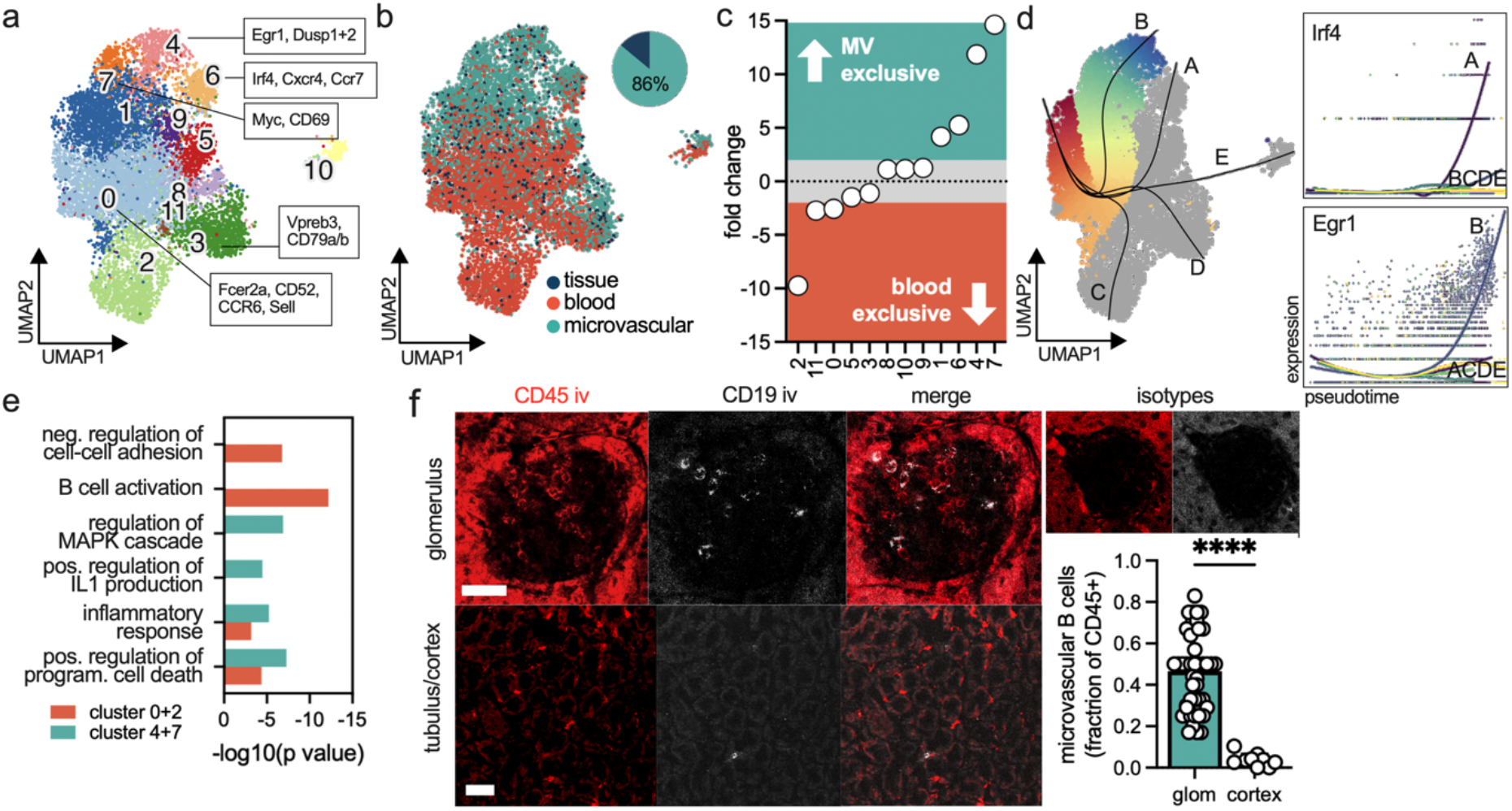
A transcriptional shift of microvascular B cells in glomerular capillaries. a) Reclustering of the B cell clusters in figure 2. Top differentially expressed (DE) genes are annotated. b) Demultiplexing of all B cells into their origins (peripheral blood, tissue or microcirculation). Of all B cells in the kidney, 86% are found in the microcirculation (pie chart). c) Ranked fold change of microvascular versus peripheral blood B cells. d) Slingshot trajectory analysis shows transcriptional phenotype switches from the main B cell cluster of the peripheral blood (cluster 0,2) to microvascular B cells of the kidney (clusters 7,4). Unique upregulation of Irf4 and Egr1 in the microvascular B cell trajectories A and B, respectively. e) Functional annotation of DE genes in peripheral blood B cells (cluster 0, 2) versus microvascular B cells of the kidney (clusters 4,7). f) Intravascular anti-CD19 B cell staining in the healthy kidney identifies the glomerular capillaries as the main effector site. Scale bar 50μm.

Next, we investigated how disease affects microvascular leukocytes. We used two different mouse models of inflammation: acute inflammation by semi-sterile peritonitis (INF) and acute kidney injury (AKI) by ischemia reperfusion (figure 4a). We hypothesized that disease remission could entail long-lasting alterations of the microvascular niche. Therefore, next to the acute phase (day 1 post surgery) we also included a late time point with complete regeneration (day 12 post surgery, INF-reg and AKI-reg, figure 4a). The kidney function was assessed by serum creatinine and renal histology, confirming injury at AKI day 1 and full renal reconstitution in AKI-reg (supplemental figure 4).

**Figure 4:**
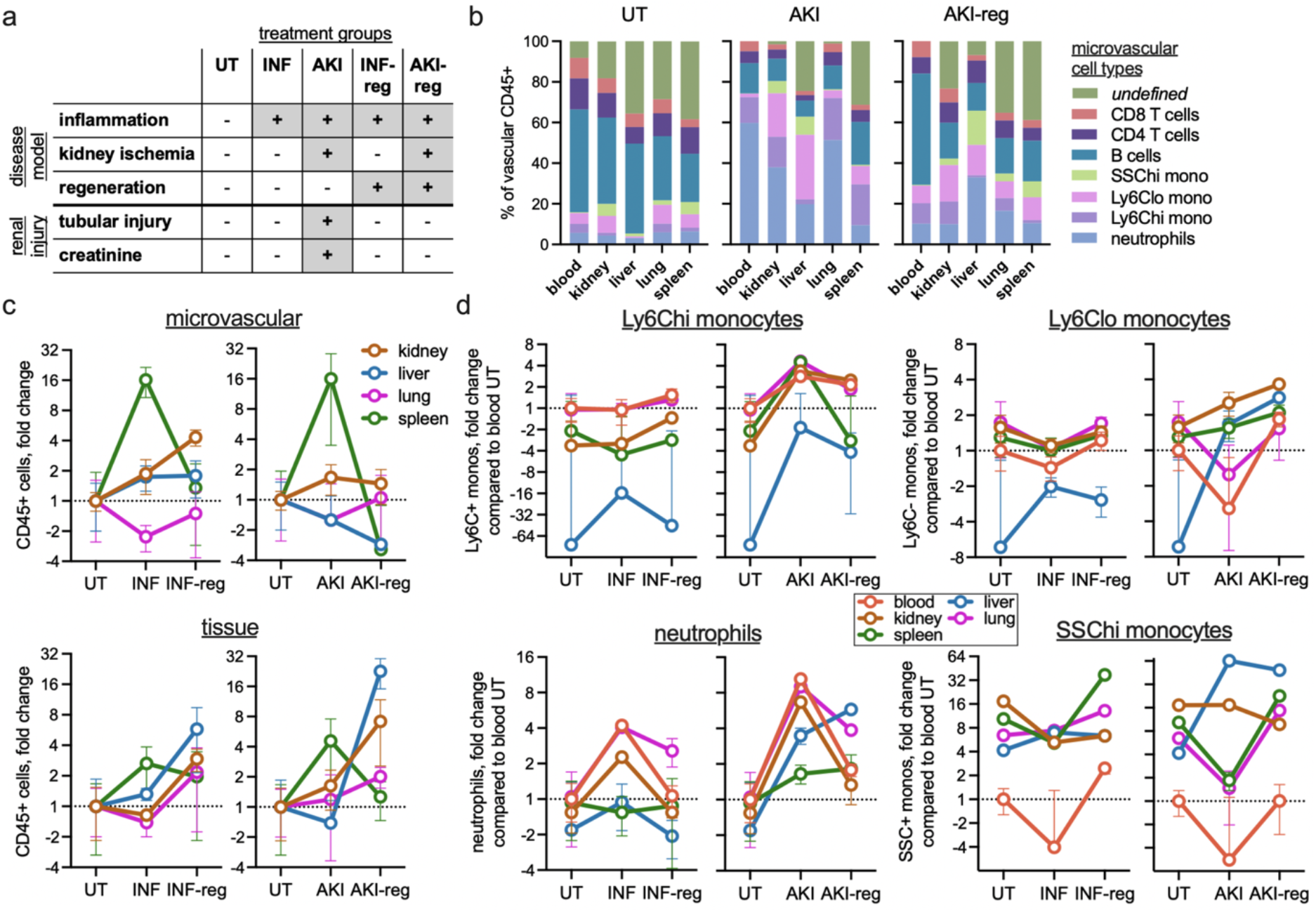
Disease-specific alterations of the microvascular immune landscape across different organs. a) Overview of the disease conditions. INF = inflammation caused by semi-sterile peritonitis. AKI = acute kidney injury. -reg = regeneration 12 days after surgery. UT = untreated. b) Microvascular leukocyte subtypes in AKI and AKI-reg compared to peripheral blood. For INF see supplemental figure 4. c) The total number of CD45+ leukocytes (microvascular = top panel, tissue = bottom panel) in selected organs expressed as relative to untreated (UT, equals 1). d) Microvascular neutrophil and monocyte subtypes in different organs and the peripheral blood expressed as relative to blood untreated (= equals 1).

**Figure 5:**
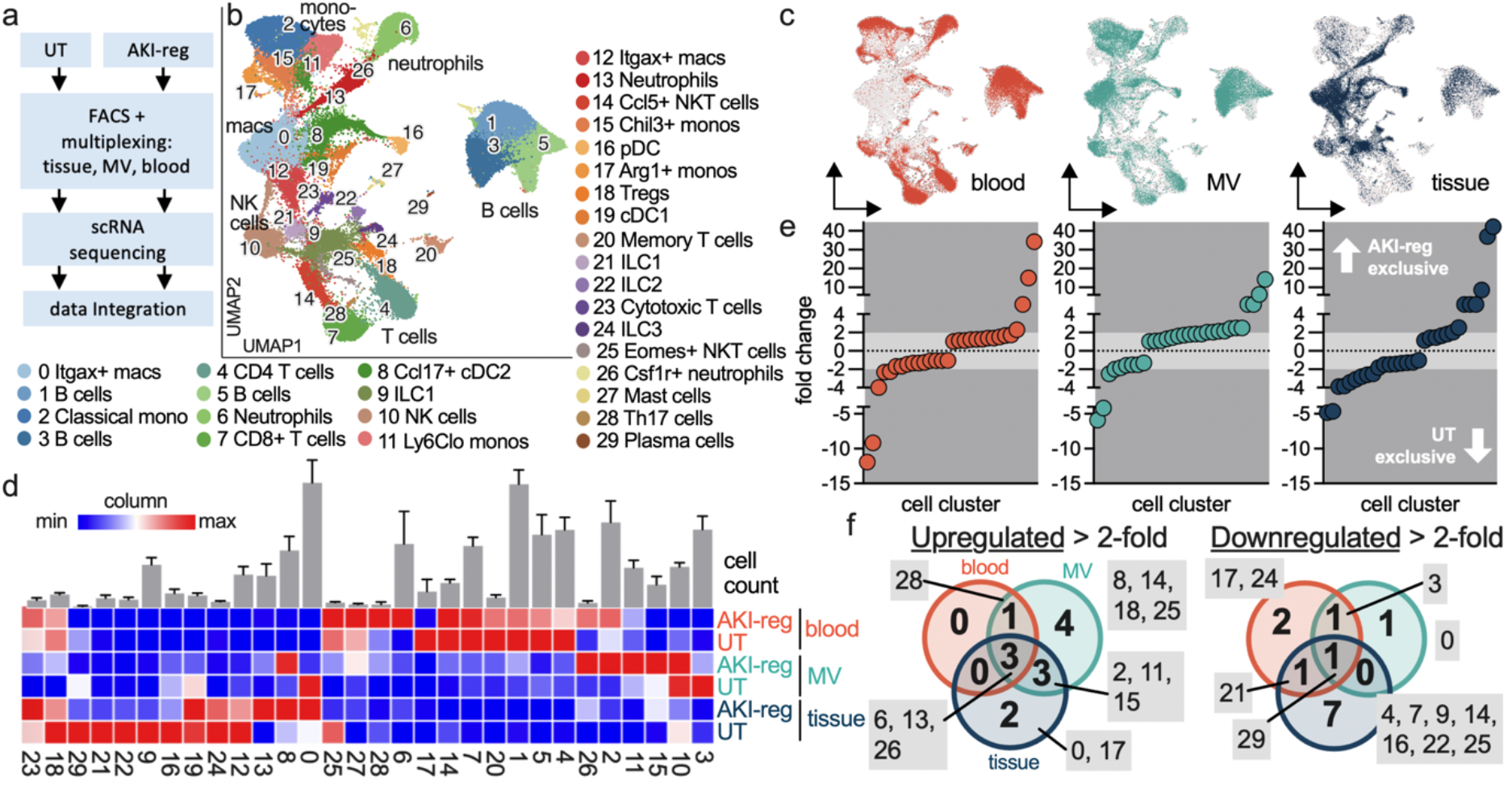
Long-term alterations of the microvascular leukocyte landscape after renal regeneration. a) Experimental setup to integrate AKI-reg (n=4) and untreated (UT, n=4) single cell sequencing data. b) Leukocyte cell clustering, UMAP visualization and subtype annotation. c) Demultiplexing, abundance of cell cluster specific leukocytes in the three compartments. d) Heatmap visualizing relative abundance of each cell cluster. Blue is the lowest number and red the highest number for each column. The top bar graphs represent the total cell count of each cluster as the mean +/- S.D. of n=8 experiments. e) Ranked fold change of each cell cluster. Negative and positive numbers mean enrichment in UT and AKI-reg condition, respectively. A detailed view is shown in supplemental figure 6. f) Up- and downregulated cell clusters in AKI-reg compared to UT. Overlapping circles indicate that the same change in a cell cluster can be detected in both (or all three) compartments. red = blood, turquoise = microvascular, blue = tissue.

We found that the total number and composition of microvascular leukocytes changed in function of time, organ and disease model, being largely independent of measurements in the peripheral blood (figure 4). During the acute phase, leukocytes increased about 16-fold in the splenic circulation compared to blood (figure 4c). In INF-reg/AKI-reg conditions, an increase of microvascular leukocytes was observed in the kidney, whereas the lung showed a reduction (figure 4c). The liver circulation was 3-fold depleted of leukocytes compared to blood in AKI-reg (figure 4c). These changes were partially uncoupled from the organ infiltration of leukocytes (figure 4c), suggesting that the vascular immune interface does not solely serve as a transmigration site. In the cell subset analysis (figure 4d), the absolute numbers were normalized to the untreated blood sample to emphasize changes in relation to blood (absolute numbers are shown in supplemental figure 5). Ly6G^+^ neutrophils sharply increased 4- to 8-fold in the blood during the acute phase in AKI and INF, and returned almost to baseline after recovery (figure 4d). However, this pattern did not reflect the neutrophil presence in the organs’ microcirculations: in the INF model, the blood neutrophil increase is not detectable in the liver and spleen microcirculation (figure 4d). In contrast to blood, neutrophils remained enriched in the lung microcirculation in both recovery models. In AKI-reg, we additionally found a specific increase of neutrophils in the hepatic circulation (figure 4d). Other examples included nonclassical monocytes and SSC^hi^ monocytes that were 16-fold and 64-fold upregulated in the liver circulation specifically in AKI-reg compared to untreated conditions (figure 4d). In these disease models, lymphoid cell types were mostly downregulated in the microvasculatures to varying degrees compared to blood (supplemental figure 4,5).

The results suggested that even full recovery from kidney injury implicates immune cell alterations at the vascular blood-kidney interface. To perform a comparative analysis, we applied single cell RNA sequencing of multiplexed leukocytes sorted from blood, the microcirculation and the tissue in AKI-reg similar to figure 2. The results were integrated with data from healthy kidneys (figure 5a). 30 leukocyte clusters could be detected based on 79.328 cells (figure 5b, supplemental figure 6). Demultiplexing assigned the cells to the specific condition (UT, AKI-reg) and anatomical compartments (tissue, blood, renal microvasculature) (figure 5c,d, supplemental figure 7). Compared to the healthy kidney, 13 leukocyte subtypes were more than 2-fold increased in AKI-reg (up to 34-fold in the blood, up to 14-fold in the renal vasculature and up to 42-fold in the tissue; figure 5e, supplemental figure 7). Of these, only four were detectable in the blood (figure 5f). Four leukocyte subtypes (clusters 14, 25, 18, 8 = NKT cells, regulatory T cells and dendritic cells) were exclusively upregulated in the renal microcirculation, three leukocyte subtypes (clusters 2, 11, 15 = different types of monocytes) were upregulated in the renal microcirculation *and* the tissue, and two leukocyte subtypes were only upregulated in the tissue (clusters 0, 17 = macrophages, monocytes; figure 5f). In conclusion, after renal recovery the majority of cellular alterations were found in the microvascular niche, none of which could be detected in the peripheral blood.

Myeloid cells are major effectors in ischemic AKI ^19^. A detailed analysis revealed different dynamics for monocytes, neutrophils and phagocytes regarding their presence in specific compartments (figure 6). Classical (cluster 2) and nonclassical (cluster 11) monocytes were present in the blood and microvasculature. In AKI-reg, we found a large increase in the renal microcirculation which was hardly reflected in the peripheral blood. Moreover, M2-like (Arg1^+^) monocytes (cluster 15) also increased but remained predominantly intravascular and not detectable in the blood (figure 6a,b). Phagocytes were absent in the peripheral blood and occupied the microvascular and tissue space to varying degrees (figure 6a,b). In AKI-reg, the largest macrophage cluster 0 lost its microvascular presence and relocated into the tissue. In contrast, cluster 8 (conventional DC type 2) expanded into the microcirculation (figure 6a,b). Neutrophils were represented by cell clusters 6, 13 and 26. All of them were strongly induced in AKI-reg (figure 6a,b). Cluster 6 predominantly defined the blood phenotype of neutrophils, whereas most microvascular and tissue counterparts belonged to cluster 13. This suggested a transformation to an infiltrative phenotype (Cxcl2^+^ Ccl3^+^ ICAM1^+^) that was already initiated in the renal vasculature (figure 6a,b). It corresponded to an activated cell state with Toll like receptor (TLR) and tumor necrosis factors (TNF) pathway enrichment (figure 6c). Consequently, the receptor-ligand analysis revealed a higher ligand density on microvascular neutrophils (cluster 13) to renal endothelium compared to blood-borne neutrophils (cluster 6) including the hypoxia-induced VEGF-A and FLT1 pathways (figure 6d) ^20^.

**Figure 6:**
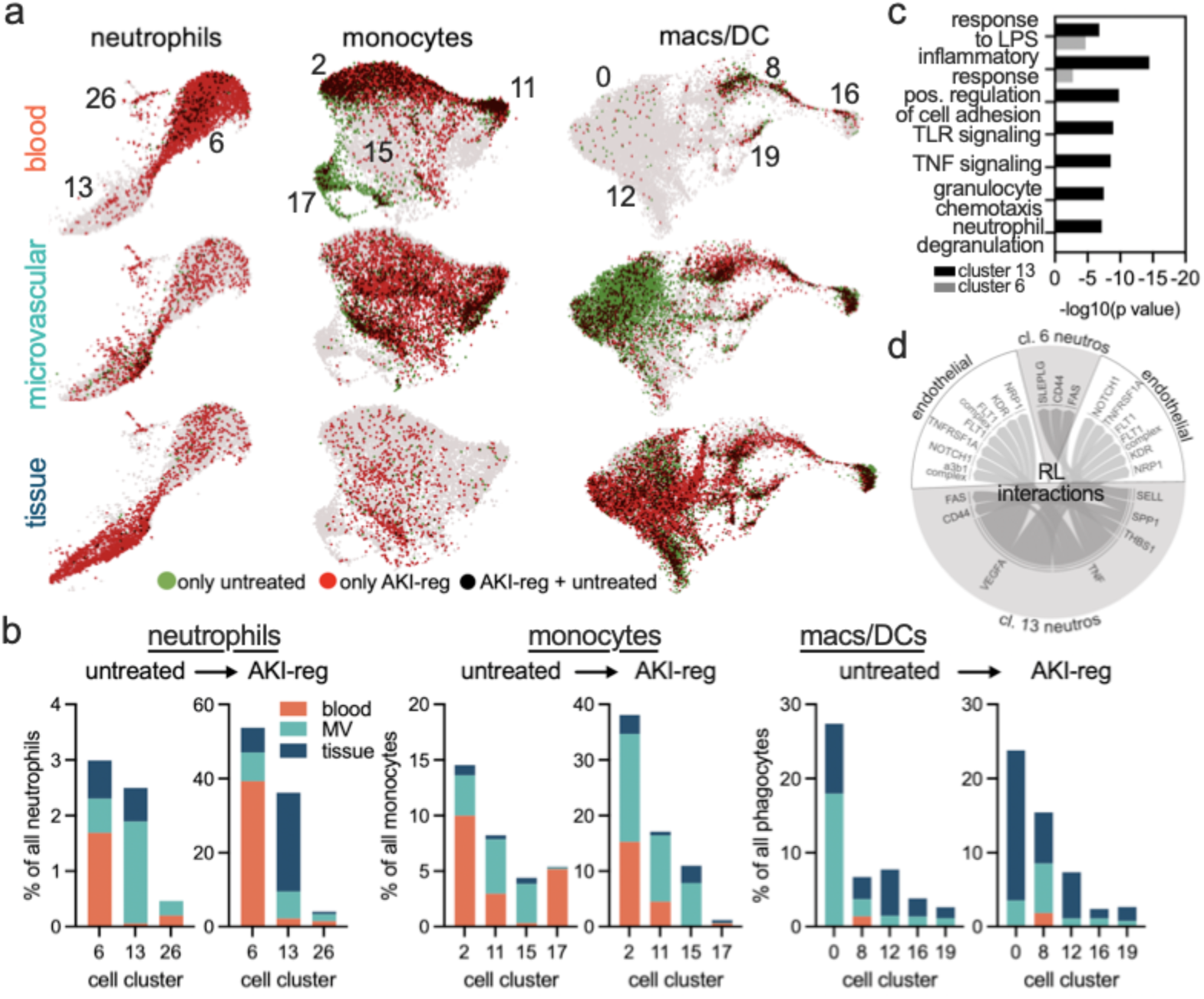
Different microvascular dynamics of myeloid cell lineages after renal regeneration. a,b) Visualization and quantification of changes in the neutrophil, monocyte and macrophage/DC lineages. Red dots = cells only present in AKI-reg conditions, green dots = cells only present in UT conditions, black dots = cells present in both conditions. c) Functional annotation of differentially expressed genes of neutrophil clusters 6 and 13. d) Predicted receptor-ligand interactions (CellPhoneDB) of neutrophil clusters 6 and 13 with renal endothelial cells.

## Discussion

Our findings highlight that the immune cell composition of blood in large veins and in the microcirculation differs significantly in an organ-dependent manner. Seven immune cell types were up to a 100-fold enriched within the healthy renal vasculature and low or virtually absent in peripheral blood. In two different disease models, numerous leukocytes showed dynamic and organ-specific alterations in the microvasculature, while no corresponding changes could be detected in the peripheral blood. Therefore, we demonstrate that microvascular immunity emerges as an inherent blind spot when relying on conventional venipuncture-based blood analysis.

The microvascular barrier comprises various cellular components, including endothelial cells, the basement membrane, and pericytes ^7,21^. Inflammation increases vascular permeability and leukocyte extravasation, making the microcirculation a focal point of injury in several diseases ^22^. Our investigation adds a new layer of structural complexity to this established concept. We have characterized a heterogeneous assembly of immune cells that are integrated into the microvascular barrier with direct contact to the blood. In the kidney, this includes leukocyte lineages such as ILC2, B cells, dendritic cells and macrophages. Some of these cells are fully ensconced within the microcirculation ^23^, while others occupy perivascular positions with intraluminal extensions ^24,25^. As exemplified by B and T cells ^26^ within glomerular capillaries, their presence can be confined to specific segments of the vascular network. These cell populations do not merely mirror the patterns of leukocyte infiltration into organ tissues. Even in inflammatory conditions, we could detect stark differences between tissue and vascular leukocytes, i.e. elevated vascular neutrophils in the liver vasculature without tissue infiltration. Thus, our findings amend the concept of the microvascular barrier by incorporating an organ-, vessel- and disease-specific cohort of leukocytes.

The presence of immune cells at the microvascular barrier with direct access to the bloodstream carries significant functional implications. Molecular signals in the plasma have the potential to trigger immediate cell activation, circumventing other regulatory components within the microvascular barrier. This finding implies a new understanding how circulating danger-associated molecular patterns (DAMPs) may mediate organ injury in various inflammatory diseases ^27^. In the context of the healthy murine kidney, the majority of intravascular leukocytes belong to DAMP-sensing phagocytes, such as conventional DC1 (Irf8^+^ Itgae^+^), plasmacytoid DC (Irf8^+^ Irf7^+^ TLR7^+^) and macrophages (Itgax^+^ Adgre1^+^ Fcgr4^+^ Tbet^+^ Gzmb^+^, and Itgax^+^ Adgre1^+^ Fcgr4^+^ Fth1^+^). These populations have been partially visualized in CX3CR1-GFP/CD11c-YFP reporter mice that feature three perivascular phagocyte populations with vascular extensions ^24,28^. As recently demonstrated by luminal monocyte aggregates in liver fibrosis ^29^, disease can fundamentally alter the vascular leukocyte composition. Our data indicate that the microvascular interface can exhibit sustained damage over an extended period of time. After regeneration following renal injury, a structural breakdown of the microvascular barrier persists, with most Fcgr4^+^ macrophages losing contact with the bloodstream and becoming concealed within the tissue. In contrast, Ccl17^+^ conventional DC2 adopt a more vascular phenotype after injury. Intriguingly, when compared to blood and renal tissue, most of the enduring changes following injury correspond to leukocyte populations within the renal vasculature. These alterations may contribute to the clinical observation that acute kidney injury elevates the overall risk of developing chronic kidney disease ^30^.

In our data set, distinct entities of the vascular tree such as arterioles, capillaries or venules cannot be distinguished. However, there are indications that microvascular leukocytes are confined to specific vascular sections. We identified a B cell subset restricted to the glomerulus. It shows a distinct vascular phenotype with a transcriptional upregulation of the maturation-associated factors Irf4 and Egr1 ^31,32^. Moreover, we revealed a substantial microvascular enrichment of ILC2. Indeed, IL5-producing ILC2s have been previously described in the perivascular space of large arteries ^33^. They are known to be responsive to various local tissue factors, including cytokines, lipid mediators, neuropeptides, and hormones ^34^. Prolonged activation through paracrine IL-33 stimulation was associated with arterial hypertrophy and eosinophilic inflammation in the lung ^35^. Our findings suggest that renal ILC2s can also respond to mediators in the blood serum, expanding our understanding of their activation mechanisms.

There are limitations to be considered. Our experimental setup only provides a snapshot in time. Thus, we cannot extract dynamic information such as directional transendothelial cell migration, intraluminal patrolling or reverse transmigration of neutrophils. Second, vessel density and intra-organ blood volumes were not accounted for. Literature reports relatively consistent values for kidney, lung, and liver in mice, suggesting that our findings are unlikely to be significantly skewed by these parameters ^36,37^. Even if vascular density slightly differed, a linear correlation between blood volume and cell numbers cannot be inferred as the hematocrit (including leukocytes), blood flow and shear stress varies along the complex microvascular vasculature ^38,39^. Thus, a direct comparison of different organs is not feasible. Finally, our microvascular cell atlas does not provide topological data. Particularly in the complex kidney architecture anatomical considerations will be important to understand the precise biological roles of different microvascular leukocytes.

Collectively, our investigations describe abundant and organ-specific leukocytes that are intimately integrated within the vascular barrier and concealed in peripheral blood. We posit that microvascular immune dynamics are critical for a more comprehensive understanding of immunity and disease perturbations.

