## Supplemental figures for "Microvascular immunity is organ-specific and concealed in peripheral blood"

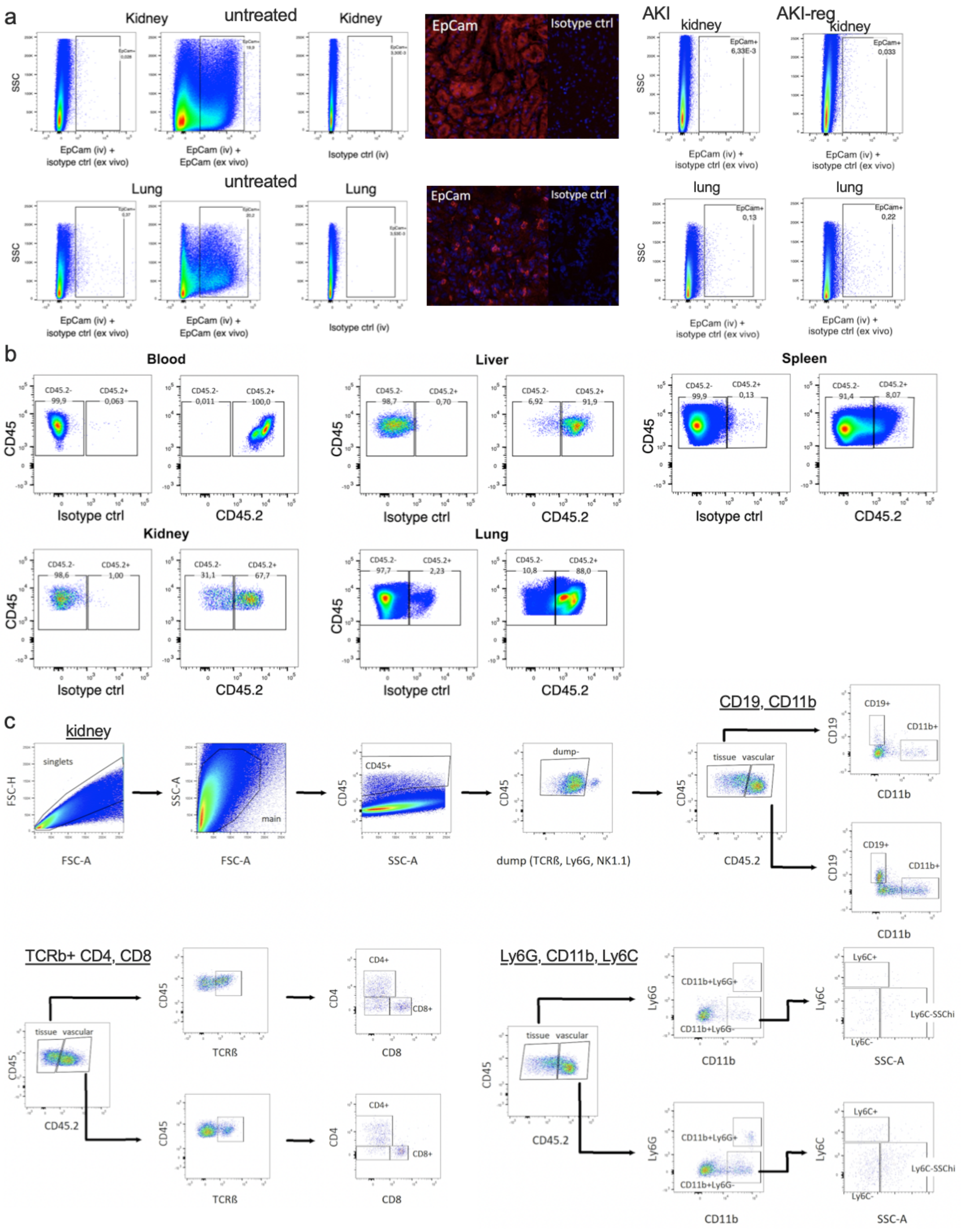


**Supplemental figure 1:** a) Intravenous injection of the anti-Epcam antibody (directed at extravascular epithelial cells) does not stain significant cell numbers in kidney and lung tissue. In contrast, the ex vivo Epcam staining detects about 20% positive cells, demonstrating that our experimental protocol specifically labels intravascular cells. b) Isotype controls of the injected anti-CD45.2 antibody for each organ. c) Gating for CD19+ B cells (top), TCRb+ CD4+ or CD8+ T cells (bottom left), CD11b+Ly6G+ neutrophils and three different types of monocytes (bottom right).

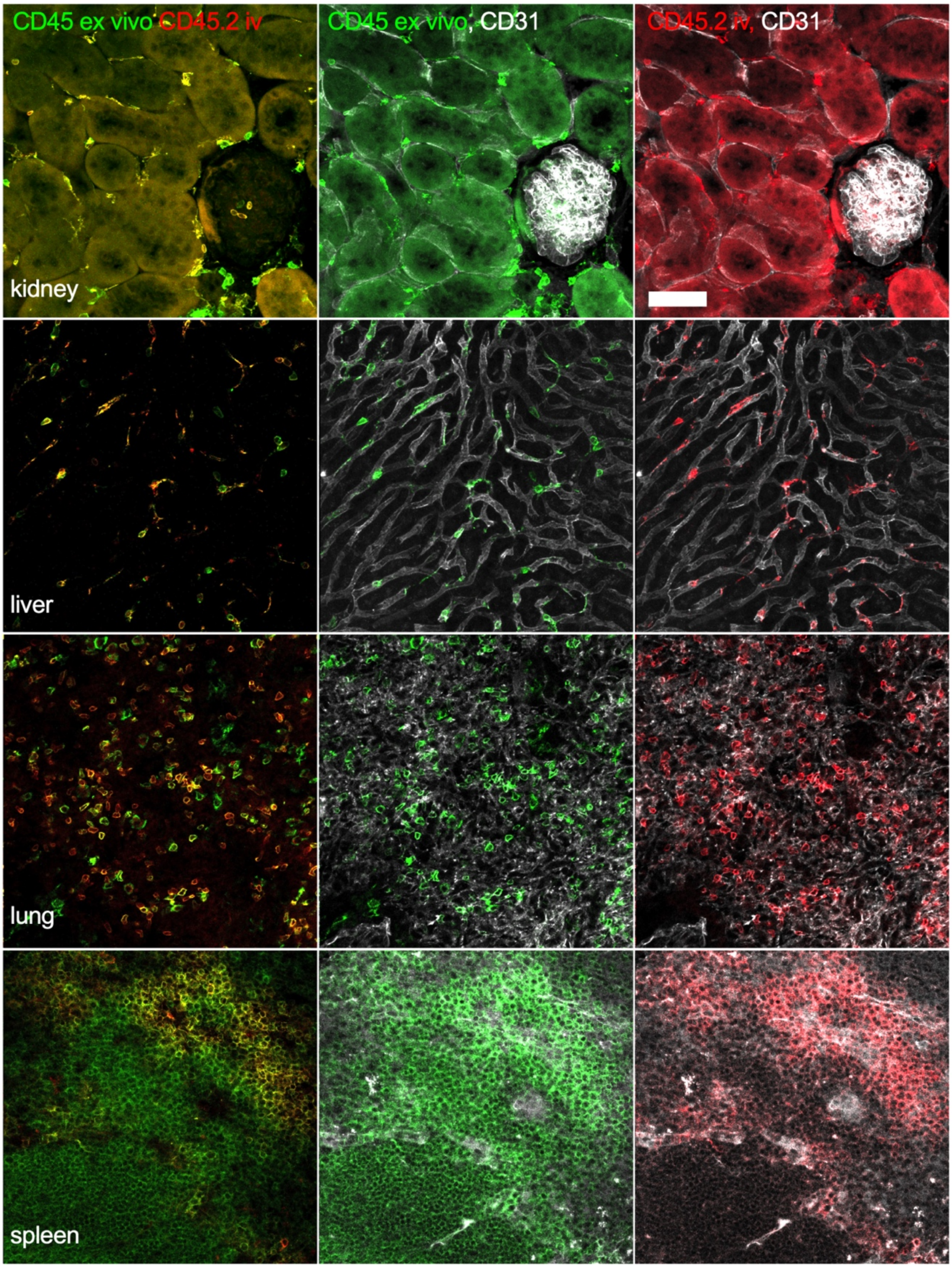


**Supplemental figure 2:** Tissue clearing and 3D confocal imaging of different organs stained with microvascular CD45.2 (red, iv injection), CD45 (green, ex vivo staining) and CD31 (white, iv injection).

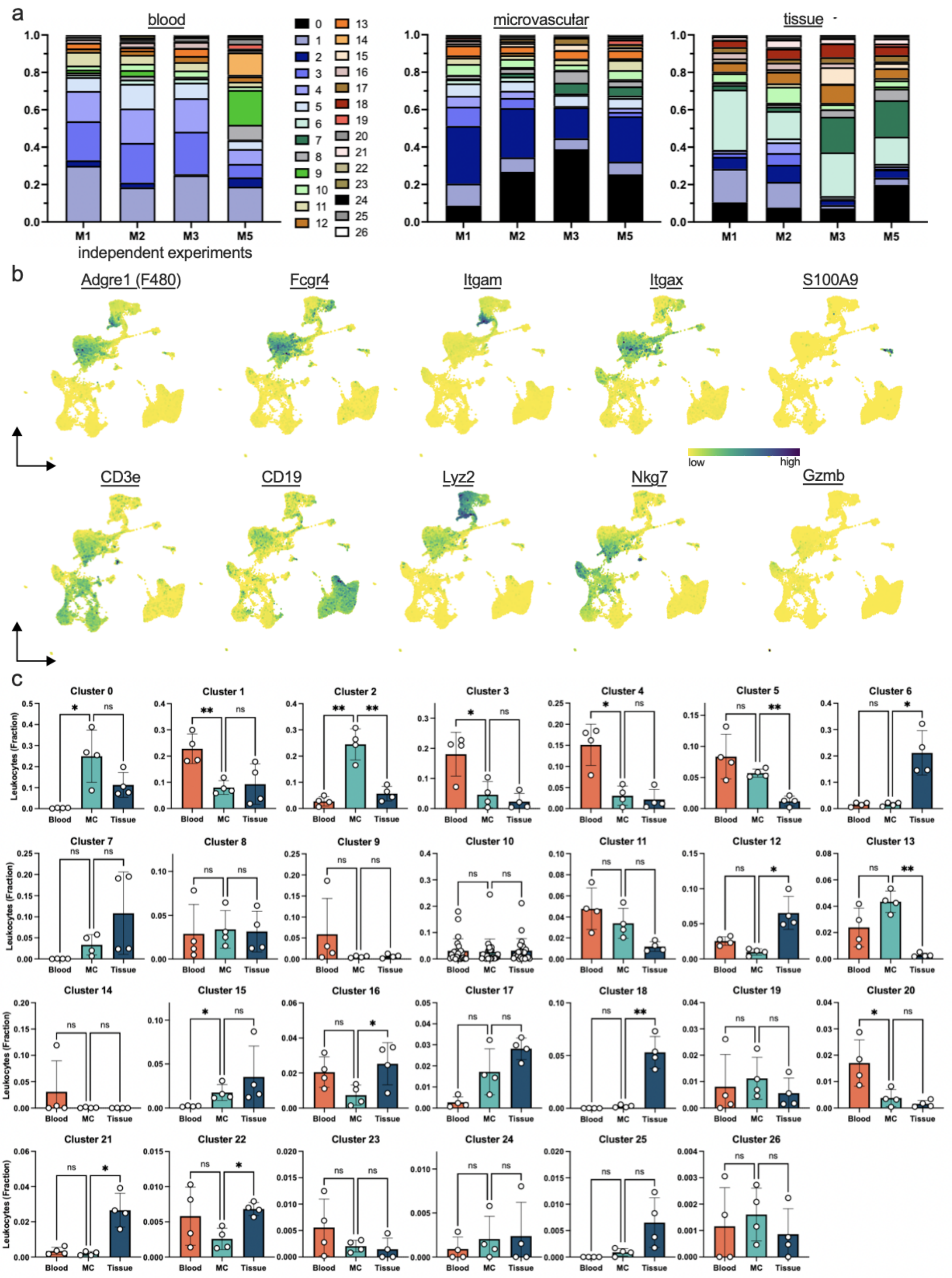


**Supplemental figure 3:** Figure corresponds to main figure 2. a) Cluster distribution of 26 cell clusters in each of the 4 untreated (UT) samples. b) Gene expression of selected genes shown on the UMAP plot. c) Leukocyte abundance for each of the 26 clusters by compartment. n=4. One-way Anova with Dunnett's correction for multiple comparison.

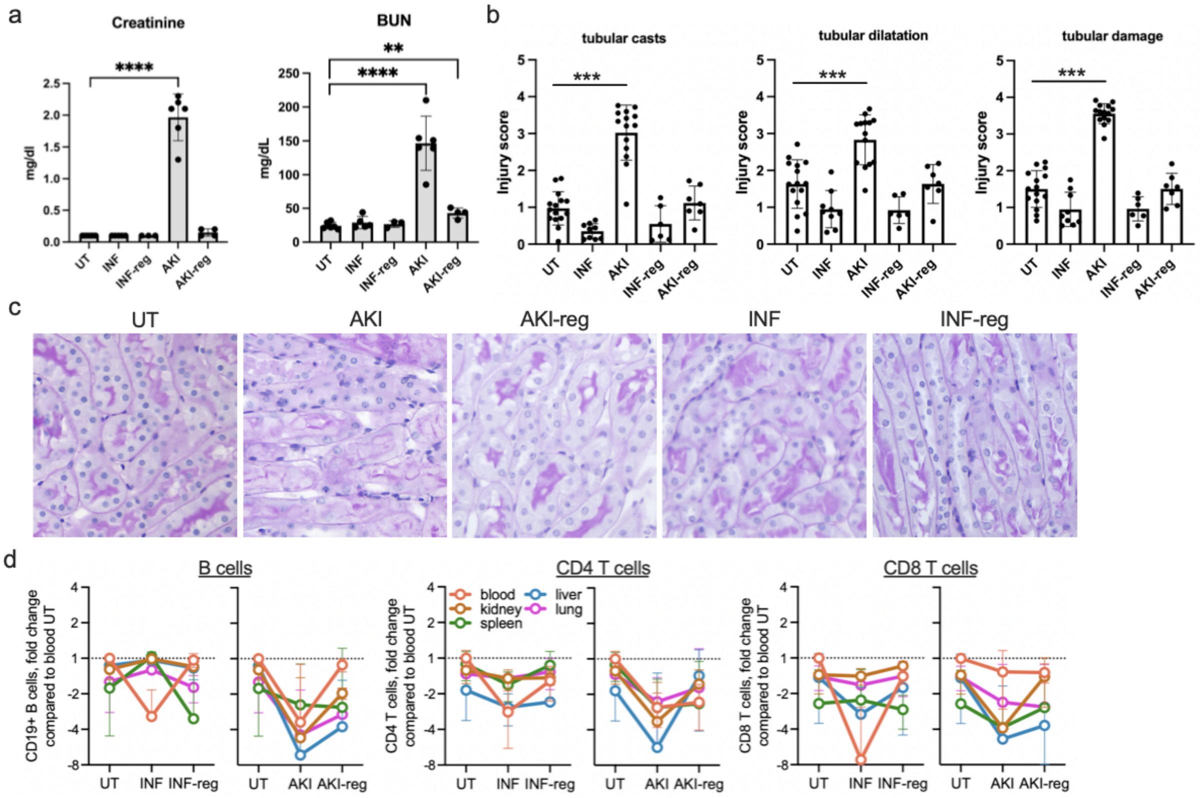


**Supplemental figure 4:** Figure corresponds to main figure 4. a) Creatinine and blood urea nitrogen (BUN) measurements in blood serum in different disease models. b,c) Histological assessment of kidney injury in different disease conditions. Representative pictures of periodic-acid Schiff (PAS) stainings. d) Microvascular lymphocyte subtypes in different organs and the peripheral blood expressed as relative to blood untreated (= equals 1).

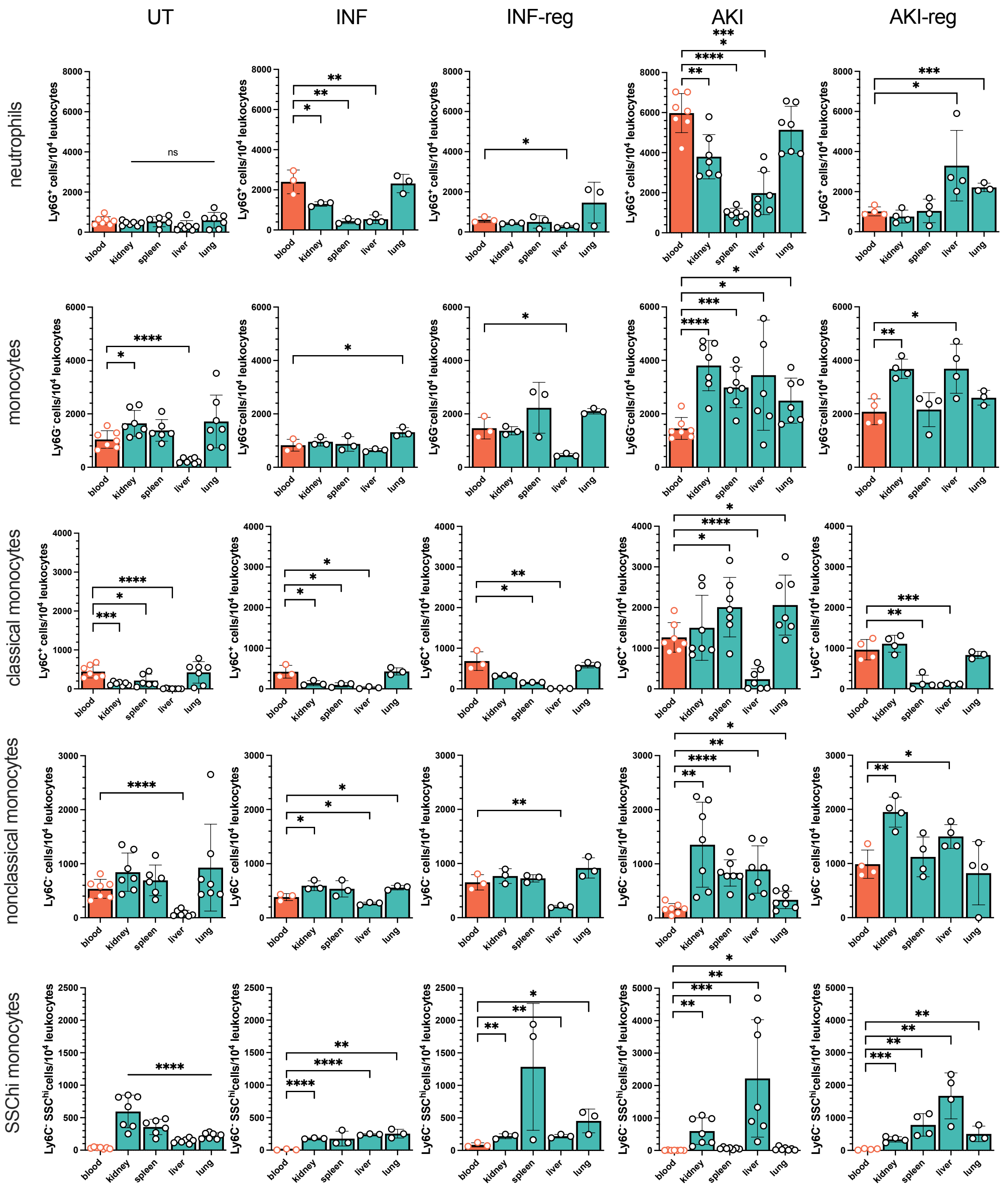


**Supplemental figure 5:** Figure corresponds to main figure 4. Flow cytometry analysis of microvascular leukocyte subtypes in comparison to peripheral blood. Unpaired t-test of blood versus each organ.

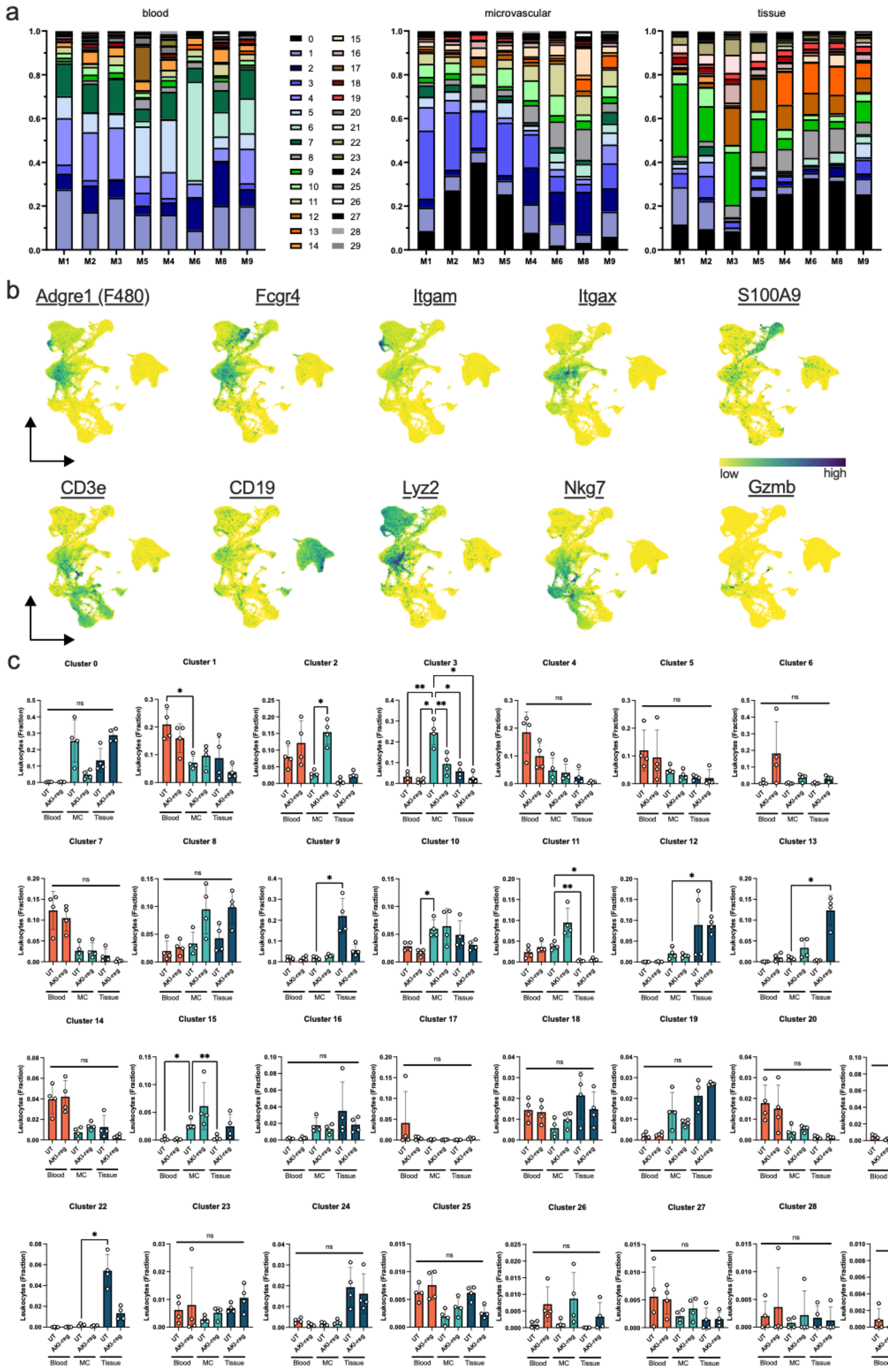


**Supplemental figure 6:** Figure corresponds to main figure 5. a) Cluster distribution of 29 cell clusters in each of the 8 samples which include 4 samples "untreated" (M1, M2, M3, M5; as shown in main figure 2) and 4 samples "AKI-reg" (M4, M6, M8, M9). b) Gene expression of selected genes shown on the UMAP plot. c) Leukocyte abundance for each of the 29 clusters by compartment. n=4 untreated and n=4 AKI-reg. One-way Anova with Dunnett’s correction for multiple comparison.

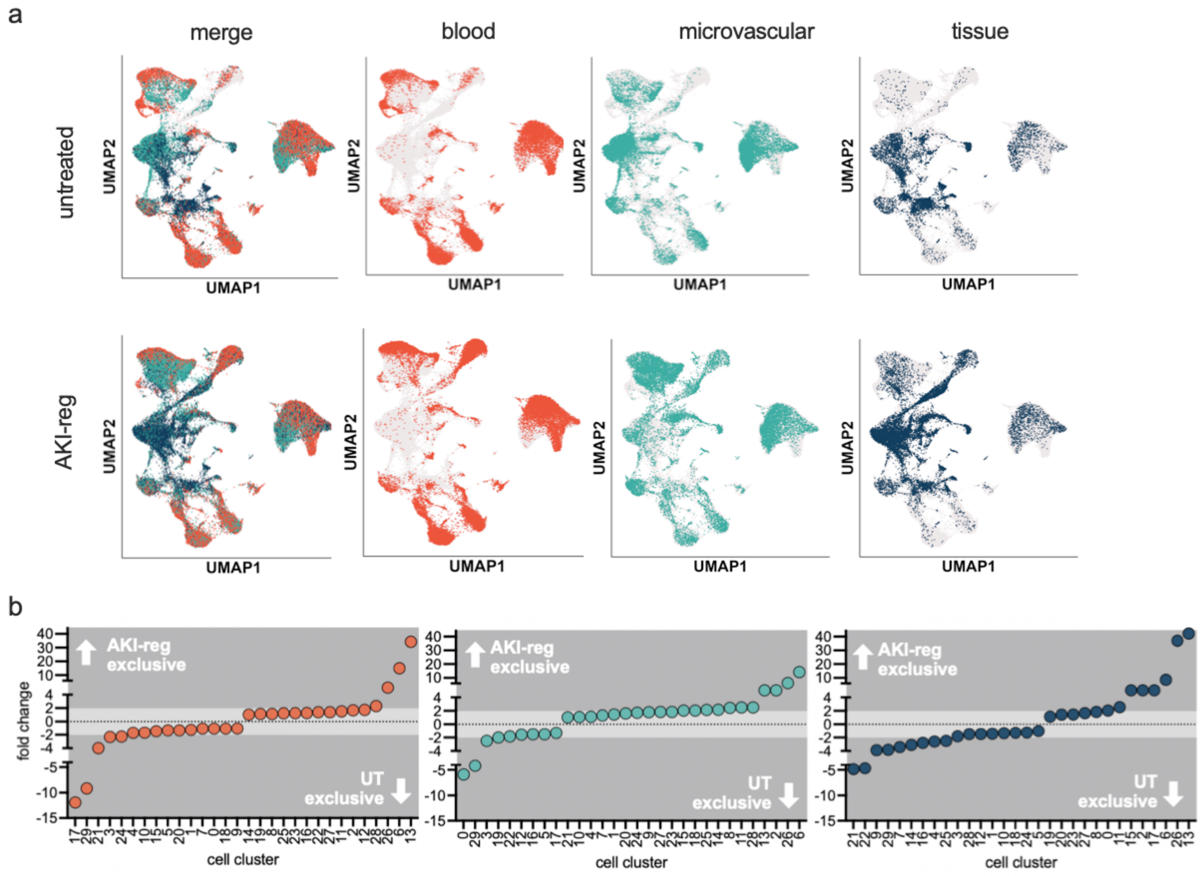


**Supplemental figure 7:** Figure corresponds to main figure 5. a) Demultiplexing into leukocytes from the blood, renal microcirculation and kidney tissue in untreated and AKI-reg conditions. b) Ranked fold change of all leukocyte cell clusters. Negative and positive numbers mean enrichment in UT and AKI-reg conditions, respectively.
